# Physical constraints on early blastomere packings

**DOI:** 10.1101/2020.05.29.123067

**Authors:** James Giammona, Otger Campàs

## Abstract

At very early embryonic stages, when embryos are composed of just a few cells, establishing the correct packing arrangements (contacts) between cells is essential for the proper development of the organism. As early as the 4-cell stage, the observed cellular packings in different species are distinct and, in many cases, differ from the equilibrium packings expected for simple adherent and deformable particles. It is unclear what are the specific roles that different physical parameters, such as the forces between blastomeres, their division times, orientation of cell division and embryonic confinement, play in the control of these packing configurations. Here we simulate the non-equilibrium dynamics of cells in early embryos and systematically study how these different parameters affect embryonic packings at the 4-cell stage. In the absence of embryo confinement, we find that cellular packings are not robust, with multiple packing configurations simultaneously possible and very sensitive to parameter changes. Our results indicate that the geometry of the embryo confinement determines the packing configurations at the 4-cell stage, removing degeneracy in the possible packing configurations and overriding division rules in most cases. Overall, these results indicate that physical confinement of the embryo is essential to robustly specify proper cellular arrangements at very early developmental stages.

**Author summary:** At the initial stages of embryogenesis, the precise arrangement of cells in the embryo is critical to ensure that each cell gets the right chemical and physical signals to guide the formation of the organism. Even when the embryo is made of only four cells, different species feature varying cellular arrangements: cells in mouse embryos arrange as a tetrahedron, in the nematode worm *C. elegans* cells make a diamond and in sea urchins cells arrange in a square configuration. How do cells in embryos of different species control their arrangements? Using computer simulations, we studied how cell divisions, physical contacts between cells and the confinement of the embryo by an eggshell affect the arrangements of cells when the embryos have only 4 cells. We find that the shape of the confining eggshell plays a key role in controlling the cell arrangements, removing unwanted arrangements and robustly specifying the proper contacts between cells. Our results highlight the important roles of embryonic confinement in establishing the proper cell-cell contacts as the embryo starts to develop.

## Introduction

During the initial stages of embryogenesis, when the number of cells (blastomeres) is very small, the spatial arrangement of blastomeres is essential for the proper development of the organism. This is particularly important in species such as ascidians, nematodes, echinoderms and mammals, whose eggs are fully divided into blastomeres (cells) upon fertilization, a process called holoblastic cleavage [1]. In embryos of these species, the spatial arrangements of blastomeres upon successive cell divisions are critical because they define the neighbors of each cell and, consequently, the signals received by each blastomere, thereby controlling cell type specification [2–5]. In nematodes (e.g., *C. elegans*) it is well established that proper contact-mediated Notch-Delta signaling between blasotmeres [5–7], which depends on the proper blastomere arrangements and their neighbor relations, is critical for the survival of the embryo. While blastomere arrangements are stereotypical for a given species, they vary substantially across species [1]. This simultaneous intraspecies robustness and interspecies variation is apparent from the early blastomere arrangements (as early as the 4-cell stage) in nematodes [8, 9], echinoderms [1, 10] and even mammals [4, 11, 12] (Fig. 1a).

**Fig 1.**
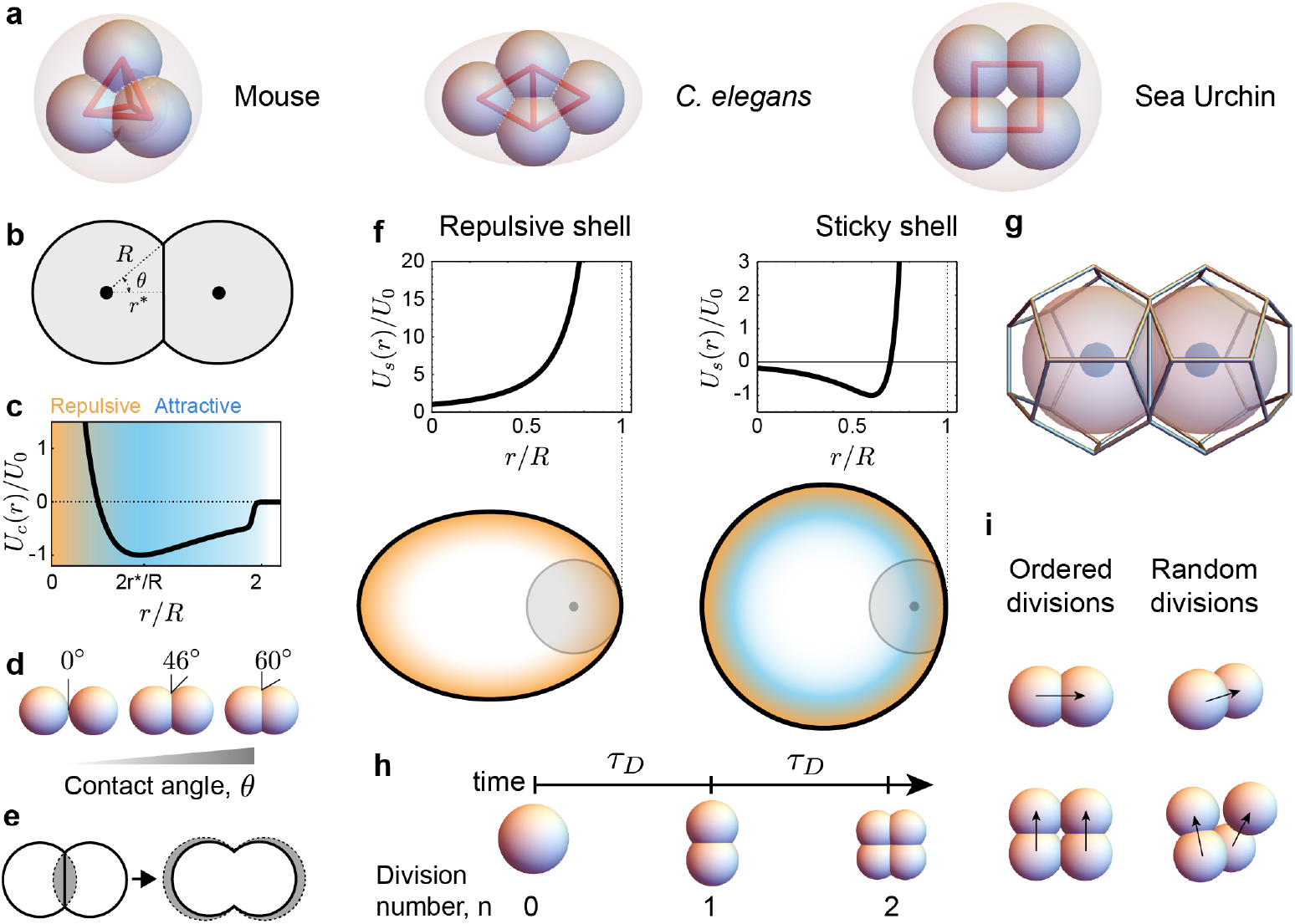
Schematics and definitions of early embryo dynamics. **a**, Schematic representation of the most common 4-cell embryo arrangements in the mouse (tetrahedron), *C. elegans* (diamond) and sea urchin (square). Blastomeres (small spheres) are confined by the surrounding confining envelope (pink; vitelline envelope, hard chitinous egg shell, or hyaline layer, respectively). **b**, Abstraction of two cells in contact, depicting the cell radius *R*, equilibrium distance *r*\* and contact angle *θ*. **c**, Cell-cell interaction potential *U_c_* (*r*). **d**, Examples of cell configurations for varying contact angles. **e**, Effective volume correction: the overlapping volume (gray, left) is added to each cell by increasing its radius to match the actual cell volume. **f**, Examples of interaction potentials, *U_s_* (*r*), of a cell with repulsive (left) and sticky (right) confining shells. Repulsive and attractive regions are shown in orange and blue tones, respectively. **g**, 3D Voronoi tessellation of neighboring cells (Methods). **h**, Definition of time between divisions *τ_D_* and division cycle *n*. **i**, Ordered divisions indicate that cells follow specific division rules. In contrast, the cell division axis is randomly oriented for random divisions.

The spatial arrangement of blastomeres in early embryos, as well as their dynamics, are ultimately controlled by their physical interactions [13]. Cell adhesion between blastomeres helps them stick together and the balance between cortical actomyosin activity and adhesion is thought to establish the contact surface between blastomeres [14–17] or, alternatively, the contact angle *θ* between them (Fig. 1b). If these were the only factors determining the arrangement of blastomeres, then the problem would be equivalent to the packing problem of a cluster of particles [18, 19], which has been extensively studied from both mathematical [20–25] and physical perspectives [19, 26, 27]. In this case, the expected cellular packing configuration (spatial blastomere arrangement) at the 4-cell stage would be a tetrahedron. While this is indeed the observed packing configuration at the 4-cell stage in mammals, the 4-cell stage packings in nematodes, ascidians, echinoderms, etc., are not tetrahedral [1]. Since the tetrahedral packing corresponds to the lowest energy state (equilibrium configuration) in particle packings, the observation of 4-cell stage packings that strongly differ from the tetrahedral arrangement indicates that either there are additional forces (beyond cell-cell interactions) affecting the blastomere equilibrium configuration, that the observed packings are metastable states with long relaxation times or that the blastomere packings are actively maintained in non-equilibrium configurations.

Beyond the direct physical interactions between blastomeres, recent experiments in *C. elegans* embryos have shown that physical confinement by the eggshell affects blastomere movements and arrangements [28–30], and several other works have highlighted the important role of division rules (i.e., the rules that define the orientation of the blastomere division planes) in blastomere arrangements [31]. The existence of cell divisions with controlled spatial orientations could maintain the system out-of-equilibrium and potentially control blastomere packings. Previous theoretical works simulating blastomere packings have either used particle-based models [28–30] or cell surface energy minimization in conjunction with a shape dependent model of division plane positioning [31–33]. However, there is no systematic study of how the different physical parameters (such as blastomere adhesion strength [34, 35] or cortical tension [15, 16]), as well as the characteristics of the confining eggshell and division rules [33, 36], affect the resulting packing configurations and their stability.

Here we sought to systematically study how the physical confinement of the early embryo, the existence of division rules and the change in adhesion/cortical tension between blastomeres control the cellular packings (blastomere arrangements) of 4-cell stage holoblastic embryos. We focus on the 4-cell stage because the observed variability across species is large, while being a tractable problem from a combinatorial and computational perspectives. By simulating the dynamics of the cells in 3D, and using Voronoi tessellation to determine the neighbor relations between blastomeres (topology of cell contacts), we find that in the absence of embryo confinement the division rules and the timing between division play an important role in the packing configurations. However, in cases for which the embryo confinement is non-negligible (as in most cases of holoblastic cleavage), the geometry of the confining shell is the main factor in the determination of the 4-cell stage cellular packings, overriding division rules.

## Methods

### Numerical integration

We solved the governing equation (Eq. 8) using the Euler-Maruyama method [37] to obtain the motion of all cells. Simulations were run using a timestep Δ*t* = 10^−3^*τ_M_*, much smaller than all relevant timescales in the system, namely *τ_M_* and *τ_D_*. The discretized version of Eq. 8 that we integrated numerically reads

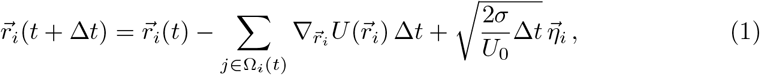

where *σ* is the magnitude of the random noise (*σ*/*U*_0_ = 5 10^−5^ for all cases), 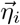 is gaussian white noise and Ω_*i*_(*t*) is the set of cells in direct contact with cell *i* at time *t*. The elements of the set Ω_*i*_(*t*) are obtained from the Voronoi tessellation of the system at time *t* (see below). All variables in Eq. 1 are normalized with their respective scales, as defined in the main text (we did not redefine the normalized variables for clarity).

Simulations were initialized with the undivided egg (first cell) at the origin for unconfined simulations or to have Gaussian distributed initial positions with variance *b*/10 around the origin for confined simulations. Simulations either ended at the timestep before cells at the 4-cell stage would divide again in non-equilibrium simulations, when 4 cells reached a tetrahedron in simulations searching for the equilibrium relaxation time, or after 8000 *τ_M_* to determine equilibrium configurations in embryos with confining shells.

### Cell-cell interaction potential

The cell-cell interaction potential has a Lennard-Jones form, but is multiplied by a support function that cuts it off at a desired distance, while keeping it continuous and differentiable (Fig 1c). The cutoff distance for a given cell pair *i* and *j* is set to *R_i_* + *R_j_* ≡ *R_ij_*, with *R_i_* and *R_j_* being the radii of cells *i* and *j*, respectively. The size of each cell (or radius, equivalently) can be different because of the volume conservation correction and because of cell divisions (see Methods below). The potential has an equilibrium distance 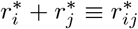 (Fig. 1b), which is the equilibrium distance between two cells combining the equilibrium radii of each cell. The specific form of the cell-cell potential *U_c_* reads

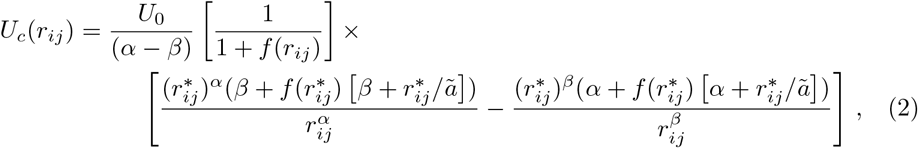

where *α*, *β*, and *a* are parameters characterizing the shape of the potential and the cutoff support function, *U*_0_ is the energy scale of the potential (the potential equals -*U*_0_ at its minimum) and 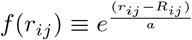. We set *α* = 4, *β* = 3, and *ã* = 0.01 in all simulations.

### Confining Shell

The shape of an axisymmetric ellipsoidal shell, and the associated ellipsoidal level set, are given implicitly by

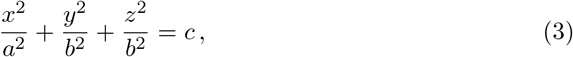

where *a* is the length of the ellipsoid’s major axis, *b* is the length of its minor axis and *c* is a positive constant that defines the ellipsoidal level set, with *c* = 1 defining the shell itself.

Since cells cannot penetrate the shell, the confining potential must diverge at the positions where the shell is located (*c* = 1). Moreover, the potential must vanish when the cell can no longer be in contact with the shell, which occurs when a cell is located at a distance larger than *R* from the shell. With this in mind, we define the confining potential of a repulsive shell *U_s_* (Fig. 1f) as

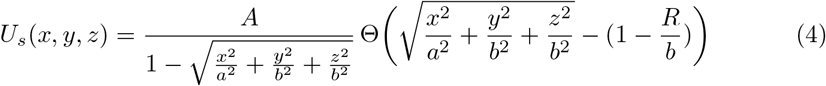

where *A* is the energy scale of the shell potential and Θ(·) is the Heaviside step function that sets the function to zero when a cell is too far from the shell to be in contact. We set *A* = 10 to balance the repulsion forces between two cells and between cells and the shell when in steady state confinement. With the interaction potential defined, the force acting on cell *i* arising from contact with the shell is given by

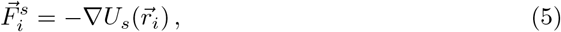

where 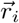 is the position of cell *i*.

In the case of a sticky shell (Fig. 1f), we use the same shell-cell interaction potential as the interaction potential between two cells, albeit with different adhesion strength. In this case, the equilibrium distance is changed to 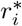 instead of 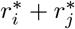 since there is only one cell interacting with the shell. In these conditions, the interaction potential for a sticky shell reads

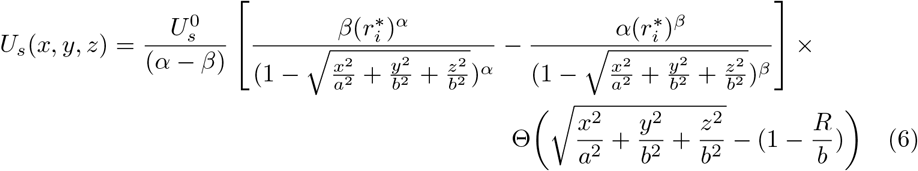

where 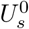 is the adhesion energy scale.

### Topology Inference

Because we simulated particles in a confined volume, it was necessary to move beyond a simple distance metric to determine if two cells were neighbors. We used a 3D Voronoi partitioning as an extra constraint in addition to distances. The package we used, Voro++ [38], determines the 3D Voronoi polytope around each cell by starting with a large 3D volume and then cutting it using the midplanes from the current cell to each of its neighbors. In this work, each cell starts with a dodecahedral volume that surrounds an inscribed sphere with the cell radius R. When a pair of cells are close enough, their dodecahedral volumes are cut by the weighted midplane between them (adjusted from the midpoint by their respective radii) (Fig. 1g). Each new cut face of the Voronoi polytope is identified with a neighboring cell allowing all neighbors to be identified. Two cells are defined to be in contact when they are within a distance *R_i_* + *R_j_* (*R_i_* and *R_j_* being the radii of cells *i* and *j*) of each other and their Voronoi polytopes share a face. An adjacency graph is created by defining each cell as a node and adding edges between each pair of cells found to be in contact. We determine the topology of each arrangement by checking if the adjacency graph is isomorphic to a reference adjacency graph for each type of topology (square, diamond, tetrahedron, T-shape, line) [22, 39].

### Cell Divisions

Cell divisions occur at well defined intervals of time *τ_D_* (Fig. 1h), with all cells dividing simultaneously. The new daughter cells were placed at a daughter radius away from the mother cell in a single timestep. In the case of random divisions, the daughter cell divides in a random direction from the mother cell (with a check to ensure that the daughter is not placed within a distance that would cause it to substantially overlap with an already existing cell). For the case of ordered divisions, the egg first divides in the x direction, then both daughter cells divide in the y direction. The total cell volume is conserved so the two daughter cells each have half the volume of the mother cell, as described in the main text.

### Volume Adjustment

The overlap between blastomeres was determined by defining a sphere with radius *R_i_* around cell *i* and then calculating its overlap volume *V_o_* with neighboring spheres. This overlapping volume then added to cell *i*, making it larger. In particular, the radius of cell *i* is modified from *R_i_* before the correction to 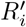 after it, with 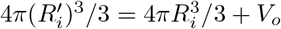 (Fig. 1e). This adjustment is performed at every timestep, and the Voronoi dodecahedron is also scaled to surround a sphere of radius 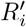 after volume correction, at each timestep too.

### Angular Mean Squared Displacement

The angular mean squared displacement is defined as

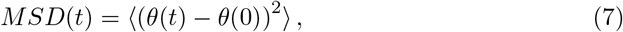

where *θ*(*t*) is the angle of a moving blastomere relative to the plane defined by three fixed cells in the x-y plane.

## Results

### Theoretical Description

In order to simulate the 3D dynamics of blastomeres, accounting for the interactions between them, their divisions as well as embryo confinement, we use a minimal representation and describe each blastomere (cell) as a particle. In this particle-based representation, cells interact with each other through an interaction potential *U*(*r_ij_*) that effectively accounts for the mechanical interactions between cells (adhesion, etc. [40]), with 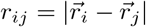 being the distance between two given cells located at positions 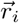 and 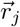. Cell-cell adhesion is represented by an attractive range in the potential, whereas a repulsive region ensures that cells do not interpenetrate when they become too close to each other (Fig. 1c). To account for cell size in this particle description, we include a sharp cut-off of the potential at a distance *R* (Methods), which corresponds to the radius of an isolated blastomere. The balance of attractive and repulsive forces between two blastomeres occurs when they are separated by a distance 2*r*\*. The ratio between this equilibrium distance between blastomeres and the blastomere size 2R corresponds to *r*\*/*R* = cos*θ*, with *θ* being the contact angle *θ* between cells (Fig. 1b). Since the contact angle is an easily measurable quantity that informs about the relative strength of adhesion and cortical tension [15, 30, 41] (Fig. 1d), we use *θ* as control parameter instead of *r*\*. Moreover, although it is not possible to enforce exact volume conservation in a particle-based description, we perform leading order corrections upon cell contact (Fig. 1e; Methods); we have checked that the volume corrections are small and we have tested that our results do not qualitatively depend on them.

At the spatial and temporal scales of embryo development, the system is overdamped and inertia can be safely neglected [42]. In this case, force balance (momentum conservation) for a given blastomere reads

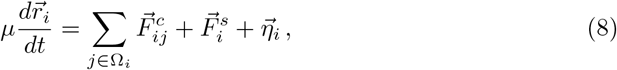

where 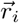 is the position of cell *i* in 3D, 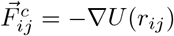 are the forces that cells in contact apply on each other (with Ω_*i*_ being the set of cells in contact with cell *i*), 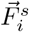 represents the force of a confining shell on cell *i* (if a confining shell is present), and 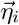 is a fluctuating force (Gaussian white noise) that is meant to represent the force fluctuations in the system (Methods). Finally, the parameter *μ* corresponds to a friction coefficient that resists cell movement in an overdamped environment and it is here assumed constant and the same for all blastomeres. To obtain the force 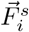 from the confining shell on cell *i*, we define the geometry of the confining shell and set the interaction potential *U*_s_(*x*,*y*,*z*) that a cell would perceive inside the shell (Fig. 1f; Methods). The confinement force perceived by cell *i* is then given by 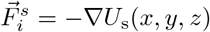.

In order to properly determine what cells are in contact and can therefore apply forces on each other, we use Voronoi tessellation (Methods; Fig. 1g). Previous particle-based simulations used distance-based metrics to determine the neighbors of each cell. However, distance-based metrics can give erroneous results for both cell-cell contacts and dynamics in the presence of confining shells. This is because when cells are highly confined, the distance between next-nearest neighbors can be smaller than the interaction potential range, thereby erroneously considering the forces of cells that are not in direct contact. Voronoi tessellation overcomes this problem and enables proper determination of cell-cell contact topology at each timestep of the simulation (Methods).

Since shells of many species have spherical or ellipsoidal shapes, we consider only these cases in what follows. We approximate the shell surrounding the embryo by an axisymmetric ellipsoid with major and minor axes *a* and *b*, respectively, with volume 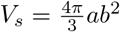 and aspect ratio *a*/*b*. Since blastomeres cannot penetrate the shell, we use confining potential forms that diverge at the shell boundary (Methods; Fig. 1f). Moreover, the confining potential vanishes for distances larger than *R* from the shell, as these distances are not within the reach of cells.

Beyond physical interactions among cells and with the confining shell, blastomeres in early embryos divide at regular intervals, with a time *τ_D_* between division events (Fig. 1h). We simulate division events accounting for the change in volume of the cells upon division (Methods). Since the volume of the daughter cells is half of cell volume before division, the cell radius *R* changes after each division cycle to *R_n_* = *R*_0_/2^*n*/3^, where *R*_0_ is the radius of the initial egg (and *V_c_* = 4*πR*_0_^3^/3 is the initial egg volume) and *n* is the number of divisions that have occurred (Fig. 1h). Finally, in order to study the role of division rules, we control the spatial direction along which cell division occurs, which corresponds to the direction perpendicular to the mitotic plane. While division rules are known to exist [33, 36, 43], the specific rules and the parameters that control them are still under debate, especially for different species. As a consequence, to study the role of division rules, we consider two limiting cases: (1) *Ordered divisions*, in which we impose representative division rules at early developmental stages (division axis is perpendicular to the division axis in the two previous division cycles; for first division, perpendicular to previous division), and (2) *Random divisions*, in which there are no division rules and we randomize the direction of cell divisions for each cell and division cycle (Fig. 1i).

Normalizing all lengths by the initial egg radius *R*_0_, all forces with *U*_0_/*R*_0_ and time with the mechanical relaxation time *τ_M_*, which is given by 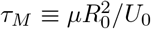 and represents the characteristic timescale over which mechanical disturbances relax to equilibrium, we obtain the relevant dimensionless parameters in the problem (Table 1).

**Table 1.** Definition of the relevant dimensionless parameters in the problem.

| Dimensionless Parameters |  |
| --- | --- |
| Parameter | Description |
| $\theta$ | *Contact angle*: Specifies the relative strength of adhesion between cells to the cortical tension of each cell. |
| $\tau_D/\tau_M$ | *Ratio of characteristic timescales*: Time between divisions relative to the mechanical relaxation time. |
| $V_s/V_c$ | *Shell to cells volume ratio*: Ratio of confining shell volume and the total volume of cells. |
| $U_s^0/U_0$ | *Cell-shell to cell-cell adhesion strength*: Ratio of the attractive energy scale for cell-shell interactions $U_s^0$ and cell-cell interactions $U_0$. |
| $a/b$ | *Shell aspect ratio*: Ratio of ellipsoid major axis length $a$ to minor axis length $b$. |

In what follows, we simulate the stochastic movements of the multiple interacting blastomeres using Langevin dynamics (Eq. 8; Methods) in different conditions.

### Unconfined Cellular Packings

To understand the packing configurations at the 4-cell stage that arise from the system dynamics, we first simulate the cellular dynamics upon divisions in the absence of embryo confinement (Fig. 2a). We define the 4-cell stage packing configurations (Fig. 2b) as the cellular arrangement just before cells at the 4-cell stage undergo the next division cycle. At equilibrium, the minimal energy configuration of 4 blastomeres in contact with each other is a tetrahedron, as already established both theoretically and experimentally for clusters of four particles with attractive interactions [21–23, 26]. However, if blastomeres divide much faster than the time required for cells to undergo mechanical relaxation (*τ_D_*/*τ_M_* < 1), cells do not have time to reach mechanical equilibrium in between divisions and the cellular packings do not coincide with the equilibrium packing configuration, as expected. When divisions are fast compared to mechanical relaxation (*τ_D_*/*τ_M_* = 0.1; Fig. 2c) and cells divide following ordered divisions, either squared or diamond configurations are observed, with diamond configurations being more prevalent as the cell contact angle increases. If cells divide in random directions, other packing configurations appear and depend on the contact angle, but both squared and tetrahedral packings are missing. In contrast to the case where division occur fast, if blastomeres take much longer than the mechanical relaxation time to divide, equilibrium packings are expected because cells should have enough time to reach mechanical equilibrium between divisions. However, our simulations show that for *τ_D_* ≫ *τ_M_* (specifically, *τ_D_*/*τ_M_* = 10; Fig. 2c), the expected tetrahedral configurations are not observed for ordered divisions (only diamond configurations are observed) and barely observed for random divisions.

**Fig 2.**
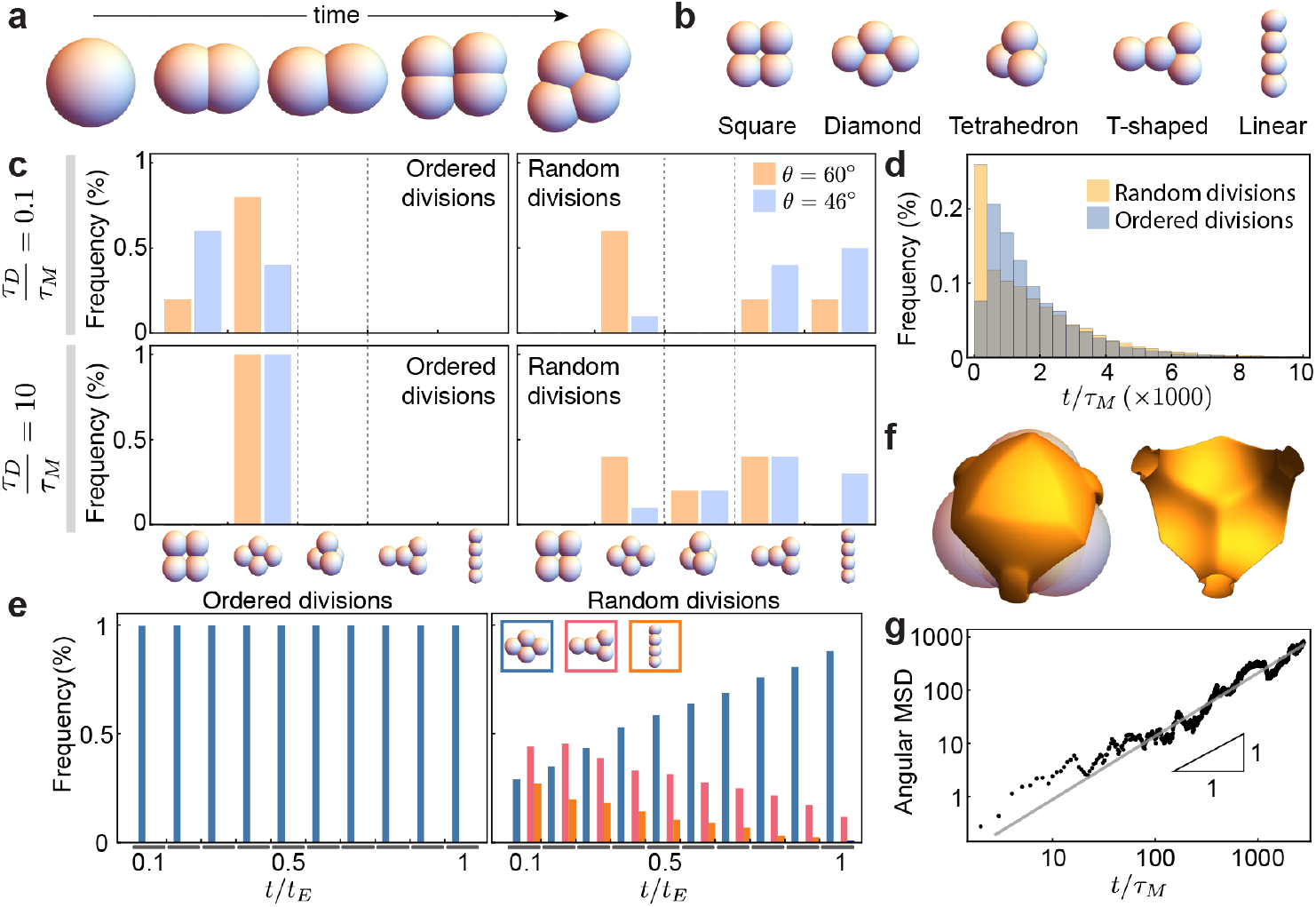
Packing configurations and dynamics of 4-cell stage unconfined embryos. **a**, Example of time evolution of cellular packings, showing how cell volume decreases by half upon division. **b**, Definition of possible topological arrangements at the 4-cell stage. **c**, Frequency of packing arrangements for slow and fast divisions (*τ_D_*/*τ_M_* = 0.1, 10, respectively), different contact angles *θ* and both ordered and random division rules (n=10^3^ simulation runs for each parameter set). **d**, Histogram of the time to reach the equilibrium tetrahedral configuration (n=10^5^ simulation runs for each condition). **e**, Frequency of packing configurations as the system relaxes to equilibrium (*t_E_* being the time to reach equilibrium) for both ordered (left) and random (right) divisions (n=10^2^ simulation runs for each condition). **f**, Top view (with cells) and cross section (without cells) showing the equipotential surface (orange tones) caused by three fixed cells on a fourth cell. **g**, Angular mean squared displacement (MSD) of a cell moving in the potential generated by three cells fixed in a triangle (n=8 simulation runs), showing its diffusive nature (fit, gray line).

To understand why the expected tetrahedral configurations are not observed, we characterized the time necessary to reach the tetrahedral equilibrium configurations at the 4-cell stage by preventing the next division round. For both ordered and random divisions, we find that cells require times three orders of magnitude longer than *τ_M_* to reach equilibrium (Fig. 2d). By monitoring the time evolution of packing configurations at the 4-cell stage as the system relaxes to equilibrium, we found that for ordered divisions cells are always in a diamond configuration before reaching the tetrahedral configuration, whereas for random divisions cells mostly evolve towards the diamond configuration and stay in that configuration until reaching the equilibrium packing (Fig. 2e). This results indicate that it takes a long time for the cluster to leave the diamond configuration, suggesting that the transition between diamond and tetrahedral configurations may involve the rotational diffusion of a blastomere. Indeed, the equipotential surface that a blastomere perceives when the three other blastomeres form a triangle indicates that in order to transit from diamond to tetrahedral configurations, one blastomere needs to traverse a flat region of the potential (Fig. 2f). The angular mean squared displacement of the movements of such blastomere scales linearly with time (Fig. 2g), showing that the transition between diamond and tetrahedral configurations occurs via rotational diffusion and explaining the long times necessary to reach the tetrahedral state.

The consequence of this floppy mode in the dynamics of the blastomeres is that it imposes an extraordinarily long time for the system to reach mechanical equilibrium, effectively leading to a degeneracy in the packing configurations at the 4-cell stage for normal division times, with degenerate packings being strongly dependent of division rules and adhesion strength between blastomeres. Such large degeneracy in the packing configurations and their strong dependency of multiple parameters is not adequate strategy to robustly specify cellular packings. Since tetrahedral packings at the 4-cell stage are observed in embryos of several species, our results suggest that another mechanism must contribute to establishing the observed tetrahedral packings, as otherwise cell divisions would need to be extraordinarily slow (*τ_D_* ∼ 24h) to allow blastomeres to reach equilibrium between divisions.

### Cellular Packings under Spherical Confinement

Many embryos displaying tetrahedral packings at the 4-cell stage feature a spherical confining shell [8, 30]. To understand the potential role of embryo confinement on cellular packings at the 4-cell stage, we simulate the dynamics of blastomeres in the presence of (repulsive) spherical confinement (Fig. 1f). We focus on long division times (*τ_D_*/*τ_M_* = 10) as this was the limit in which tetrahedral packings were expected, but shown above to be missing due to the long times associated with rotational diffusion of the blastomeres. If the confining shell has a very large volume compared to the total volume of the cells (*V_s_*/*V_c_* ≫ 1; Fig. 3a), the situation is similar to the unconstrained embryo. As the volume of the confining shell is decreased, the 4-cell stage packings start to change because cells start interacting with the repulsive shell. Finally, when the volume of the confining shell is comparable to the volume occupied by the cells (*V_s_*/*V_c_* ≃ 2), only tetrahedral configurations are observed (Fig. 3a). In this case, we find that blastomeres robustly reach the tetrahedral packing at the 4-cell stage five orders of magnitude faster than in the absence of confinement (relaxation time < *τ_M_*; Fig. 3b) and irrespective of their contact angle or division rules (Fig. 3c). These results indicate that the presence of spherical confinement removes the degeneracy of cellular packings and quickly imposes a tetrahedral blastomere configuration, overriding division rules.

**Fig 3.**
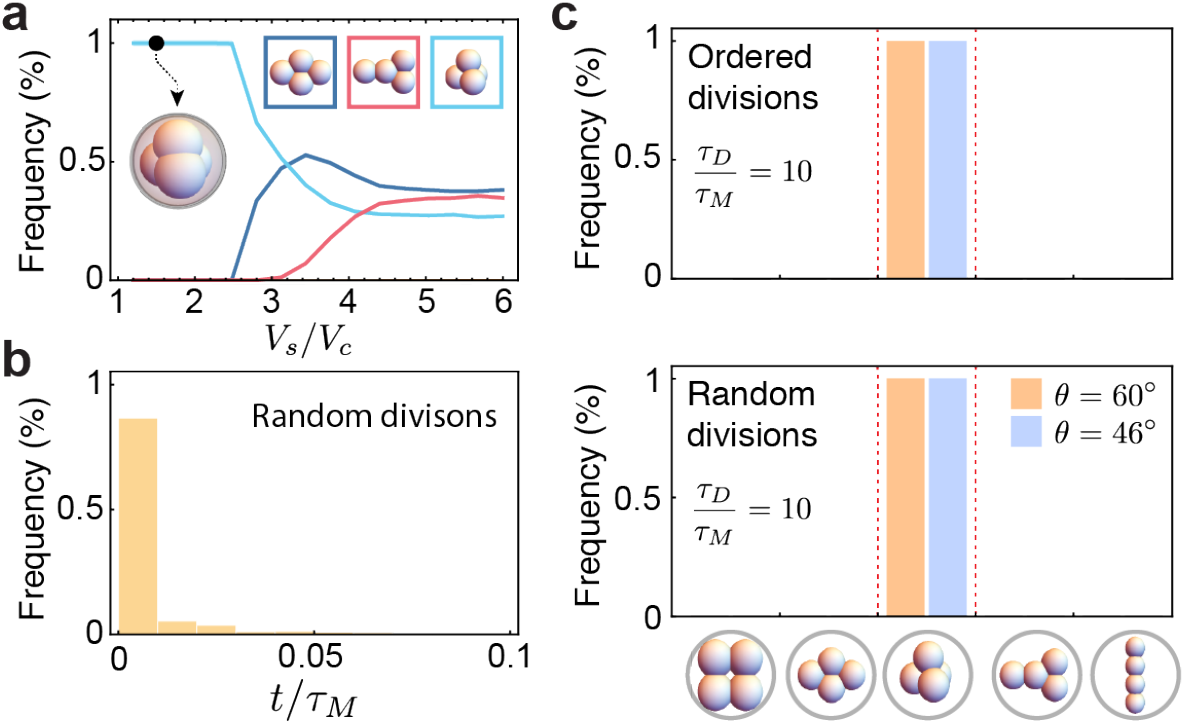
Spherically confined cellular packings. **a**, Frequency of different cell arrangements for embryos with spherical repulsive confinement as the ratio of the shell volume and total cells volume varies (random divisions; *τ_D_*/*τ_M_* = 10; n=10^4^ simulation runs for each parameter set). **b**, Histogram of the time to reach the equilibrium tetrahedral configuration (n=10^3^ simulation runs). **c**, Frequency of packing arrangements for embryos confined in a spherical repulsive shell (*a*/*b* = 1; *V_s_* /*V_c_* = 1.52), and both slow and fast divisions (*τ_D_*/*τ_M_* = 0.1, 10, respectively), different contact angles *θ* and both ordered and random division rules (n=500 simulation runs for each parameter set).

### Cellular Packings under Ellipsoidal Confinement

While embryos of several species have spherical confining shells, other shell geometries are observed across species. Different nematode species display elongated axisymmetric shells of varying aspect ratios [8, 9] that can be approximated by an axisymmetric ellipsoidal geometry. Previous works have shown that the shape of the confining shell is important for cellular arrangements in nematodes [30], for which the 4-cell stage packing arrangements are critical for the survival of the embryo, as improper cell contacts lead to fatal developmental defects [44]. To understand the role of varying ellipsoidal shell geometries on the cellular packings across nematode species, we systematically studied the effect of confining shell volume and aspect ratio on the blastomere packing configurations at the 4-cell stage.

Similarly to spherical shell geometries, for all simulated aspect ratios of ellipsoidal shells (*a/b* = 1,…, 3) we find that when the confinement is negligible (*V_s_*/*V_c_* ≫ 1) and divisions are randomly oriented, packing configurations are strongly degenerate even if the time between divisions is much longer than the mechanical relaxation timescale (*τ_D_*/*τ_M_* ≫ 1; Fig. 4). In this case, different packing configurations have similar relative frequencies, albeit with the diamond configuration always being predominant. Essentially, if the volume of the shell is sufficiently large compared to the total volume of the blastomeres (*V_s_*/*V_c_* ≫ 1), the observed packings at the 4-cell stage are similar to unconfined embryos (Figs. 2c and 4), as expected. As the volume of the shell is decreased and the blastomeres start to feel the physical confinement, the relative frequencies of 4-cell stage packings start to change, removing some degeneracy in packing configurations, in a manner that depends on the shell aspect ratio. When the volume of the confining shell is comparable to the volume occupied by the cells (*V_s_* ≃ 2 *V_c_*), the degeneracy in 4-cell stage packings largely disappears and different, but unique, packings exist for different aspect ratios (Fig. 4).

**Fig 4.**
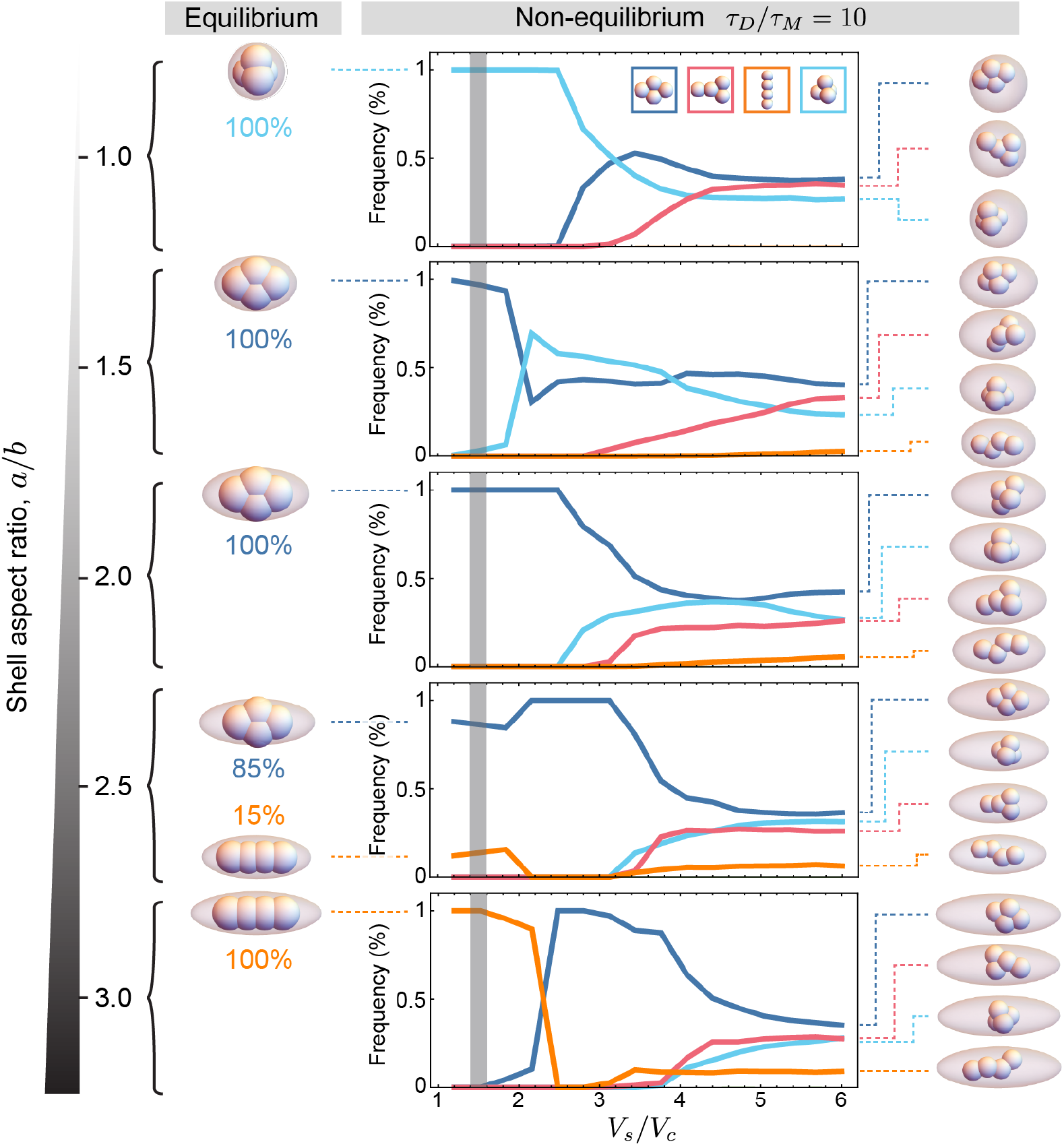
Cell arrangements for repulsive ellipsoidal confinement. Frequency of different cell arrangements for embryos with ellipsoidal repulsive confinement of varying aspect ratio (*a/b*) as the ratio of the shell volume and total cells volume varies (random divisions; *τ_D_* /*τ_M_* = 10; n=10^4^ simulation runs for each parameter set). Snapshots of non-equilibrium packing configurations for large shell volumes (*V_s_* /*V_c_* = 6) are shown on the right. Equilibrium (*t* = 8000*τ_M_*) packing configurations for each aspect ratio and *V_s_*/*V_c_* = 1.52 are shown on the left (n=200 simulation runs for each parameter set).

For some shell geometries we observe that even under confinement, two different packing geometries are possible, albeit with different relative frequencies (Fig. 4). For aspect ratio of 1 (spherical limit), only tetrahedral packings are obtained, as described above and observed for nematode species with spherical shells [8, 30]. As the aspect ratio increases the relative frequency of the tetrahedral packing diminishes and the frequency of diamond packings increases, with only diamond configurations observed between *a/b* ∼ 1.5 and *a/b* ∼ 2.4. Increasing the aspect ratio above *a/b* ∼ 2.4 leads to the coexistence of diamond and linear configurations (Fig. 4; *a/b* ∼ 2.5). For aspect ratios of a/b ∼ 3 and above, the only observed configuration is linear.

To check if the packing configurations obtained in confined embryos (*V_s_*/*V_c_* = 1.52; *τ_D_*/*τ_R_* = 10) correspond to the actual equilibrium packings for each specific shell geometry, as was the case for the spherical shell, we simulate the dynamics of blastomeres, preventing further cells divisions at the 4-cell stage and letting the system reach equilibrium. We find that for each value of the aspect ratio, the packing configurations observed for *τ_D_*/*τ_R_* = 10 (with random divisions) were the actual equilibrium configurations of the blastomeres at the same confining volume (*V_s_* /*V_c_* = 1.52) and aspect ratio. This indicates that the confining shell eliminates the degeneracy in packings and selects the equilibrium packing configurations for a given shell geometry. These results indicate that the geometry of the confining shell alone can determine the 4-cell stage blastomere arrangements regardless of the specific division rules, providing a robust mechanism to remove packing degeneracy and select the proper cellular packing.

### Cellular packings in sticky shells

So far, we have only considered shells that confined the blastomeres by generating a repulsion force upon contact (repulsive shell, Fig. 1f). However, in some species, the blastomeres can adhere to the confining shell, likely affecting blastomere packing configurations. In the case of sea urchin embryos (echinoderms), there is evidence of strong adhesion to the hyaline layer surrounding the blastomeres [10, 45–47] and the 4-cell stage blastomere packing configuration is a square [1], a configuration never observed in the cases described above. In the case of sea urchins, the geometry of the hyaline layer (confining shell) that surrounds the blastomeres is not exactly spherical and changes slightly over time. However, for the sake of simplicity, here we consider a spherical sticky confining shell (Fig. 1f). Since echinoderms have stereotypical division rules (dividing perpendicular to the two previous divisions at early stages) we study the effect of shell-blastomere adhesion strength and confining volume on the 4-cell stage packing configurations for ordered divisions (Fig. 1i).

For large shell volume compared to the total blastomere volume (*V_s_*/*V_c_* ≫ 1), the only observed configuration with finite blastomere-shell adhesion is the diamond configuration, with all blastomeres adhered to the shell (Fig. 5a). When the confining volume becomes comparable to the blastomeres volume (*V_s_*/*V_c_* ≃ 1 – 1.5), the diamond configuration is suppressed and the tetrahedral and square packing configurations coexist at different frequencies depending on the relative strength of cell-cell adhesion and cell-shell adhesion (Fig. 5b,c). In this case, if adhesion to the shell is very low, only tetrahedral configurations are observed, as expected in the limit of negligible shell adhesion (repulsive shell). When the adhesion to the shell dominates over cell-cell adhesion, we find that square and tetrahedral packing configurations are observed approximately 40% and 60% of the time, respectively.

**Fig 5.**
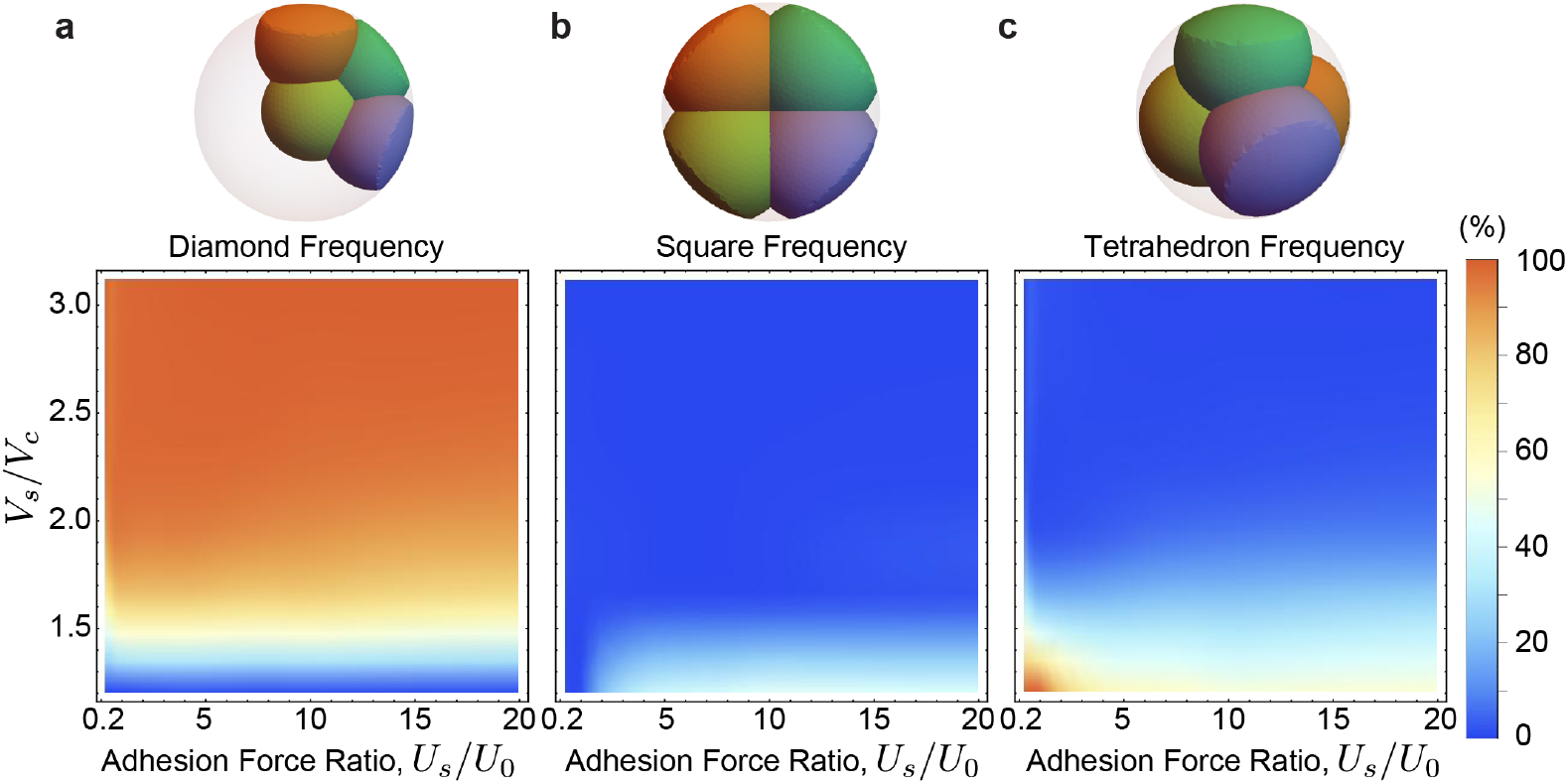
Cell packing arrangements in a sticky confining shell. **a-c**, Frequency of packing arrangements (a, diamond; b, square; c, tetrahedron) for varying values of the shell volume to cells volume ratio, *V_s_*/*V_c_*, and cell-shell to cell-cell adhesion strength, 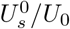 (n=5000 simulation runs for each parameter set).

The existence of strong adhesion to the confining shell introduces square packing configurations that were not observed in purely repulsive confining shells. However, the blastomere adhesion to the shell cannot be the only reason why 4-cell stage embryos of echinoderm species are square, since there is still a strong probability of tetrahedral packings even in the presence of adhesion.

## Discussion

Proper blastomere arrangements in early embryogenesis, and in particular their topology of cell-cell contacts, are critical to ensure proper development. Here we presented a systematic study of the possible (non-equilibrium and equilibrium) packing configurations (cell arrangements) both in the absence and presence of a confining shell that physically restricts the movements of blastomeres. We find that the shape of the confining shell determines the blastomere packing configurations of 4-cell stage embryos, regardless of division rules, removing blastomere packing degeneracies that could lead to fatal developmental defects.

In the absence of a confining shell, we find that the relaxation time to reach the equilibrium configuration is extraordinarily long due to rotational diffusion of blastomeres. Fast blastomere divisions generate a considerable degeneracy of 4-cell stage packings, which are sensitive to adhesion levels, division times and division rules. In this scenario, a very tight control of division axis and timings would be necessary to ensure proper embryonic packings. While it would be possible to find a set of division rules and timing of divisions to encode virtually any blastomere packings, in this scenario the packing configurations would be highly sensitive to noise and not very robust. In embryos with not confining shell, or in meroblastic cleavage, the attachment of cells to the yolk may prevent slow rotational diffusion of blastomeres. In this case, the division rules [33, 36] are essential to control blastomere packing configurations [31].

Our results indicate that the presence of a confining shell removes degeneracies in the 4-cell stage packing configurations and robustly establishes a stable configuration, solely dependent on the shape of the confining shell. For spherical shells, this leads to a unique tetrahedral packing, as observed in mouse embryos [4, 12] and nematode worms with spherical confinement [8, 9, 30]. Our results also reproduce the observed packing configurations in different species of nematodes with varying degrees of shell elongation [8, 30], and are in agreement with previous models and experiments of this process [30]. The presence of the shell also decreases significantly the time for the blastomeres to reach the equilibrium configuration. In this case, division rules could only control the packings if blastomeres divided extraordinarily fast (< 0.1 min; *τ_M_* ∼ 1 min), an unlikely scenario. In essence, the role of the confining shell is to enforce a robust 4-cell stage packing configuration that depends only on the shell geometry and is largely insensitive to division rules or noise in the system.

In the case of sticky confining shells, our work suggests that the robust square arrangement observed in echinoderms cannot be explained solely by the strong adhesion to a spherical shell (hyaline layer), since the experimentally observed square blastomere arrangement was only obtained approximately 50% of the times in the simulations. Observations of the hyaline layer in the early sea urchin embryo show that it closely surrounds the blastomeres and that it plastically deforms upon divisions. The precise shape of the hyaline layer and its temporal changes were not accounted for in our simulations and are likely to play an important role in determining the 4-cell stage blastomere packings.

Unlike previous particle-based descriptions [30], we introduced 3D Voronoi tessellation to determine the topological blastomere arrangements (cell contacts). Using a distance metric to obtain topologies can lead to erroneous contacts and unphysical dynamics. Instead, tracking the system topology with a 3D Voronoi tessellation allows the correct calculation of the forces between cells even for high contact angles or for cells under strong confinement. However, as in any particle-based model, our description does not account for the effect of changes in cell shape on the resulting force between cells or contacts, which may be relevant in some situations to accurately predict blastomere motions or the axis of cell divisions. Previous particle-based models have shown that it is possible to properly account for the dynamics of the blastomeres in early *C. elegans* embryos [29]. However, it is unclear if in some cases where cell shape changes are different and more complex, particle-based models would still be accurate. In contrast to particle based models, other descriptions simulate the cells shapes to account for geometry-dependent division rules [31]. These descriptions rely on energy minimization (Surface Evolver [48]) to obtain cell shapes and are therefore limited to equilibrium packings. Our work combines the fast simulation power of particle description with Voronoi tessellation to determine cell neighbors (contact topology). We expect our description to fail if cell shapes are not compact (e.g., very elongated cells) because in these conditions the Voronoi tessellation would not provide a faithful representation of cell-cell contacts. Using this hybrid simulation method, we can explore the non-equilibrium dynamics of cellular packings that cannot be captured by equilibrium descriptions.

Altogether, our work demonstrates that physical confinement provides a powerful way of robustly guiding the blastomeres to one particular arrangement, strongly reducing the set of possible arrangements an embryo could take and helping guide the embryo developmental trajectory.

## Acknowledgments

We thank all Campas lab members for thoughtful discussions and input. We acknowledge support from the Center for Scientific Computing from the CNSI, MRL: an NSF MRSEC (DMR-1720256) and NSF CNS-1725797.

